# BMP signalling facilitates transit amplification in the developing chick and human cerebellum

**DOI:** 10.1101/2020.10.12.335612

**Authors:** V Rook, P Haldipur, K Millen, T Butts, RJ Wingate

## Abstract

The external granule layer (EGL) is a transient proliferative layer that gives rise to cerebellar granule cell neurons. Extensive EGL proliferation characterises the foliated structure of amniote cerebella, but the factors that regulate EGL formation, amplification within it, and differentiation from it, are incompletely understood. Here, we characterise bone morphogenic protein (BMP) signalling during cerebellar development in chick and human and show that while in chick BMP signalling correlates with external granule layer formation, in humans BMP signalling is maintained throughout the external granule layer after the onset of foliation. We also show via Immunohistochemical labelling of phosphorylated Smad1/5/9 the comparative spatiotemporal activity of BMP signalling in chick and human. Using *in-ovo* electroporation in chick, we demonstrate that BMP signalling is necessary for subpial migration of granule cell precursors and hence the formation of the external granule layer (EGL) prior to transit amplification. However, altering BMP signalling does not block the formation of mature granule neurons but significantly disrupts that pattern of morphological transitions that accompany transit amplification. Our results elucidate two key, temporally distinct roles for BMP signalling *in vivo* in organising first the assembly of the EGL from the rhombic lip and subsequently the tempo of granule neuron production within the EGL.

**Significance statement:** Improper development of cerebellar granule neurons can manifest in a plethora of neurodevelopmental disorders, including but not limited to medulloblastoma and autism. Many studies have sought to understand the role of developmental signalling pathways in granule cell neurogenesis, using genetic manipulation in transgenic mice. To complement these insights, we have used comparative assessment of BMP signalling during development in chick and human embryos and *in vivo* manipulation of the chick to understand and segregate the spatiotemporal roles of BMP signalling, yielding important insights on evolution and in consideration of future therapeutic avenues that target BMP signalling.

## INTRODUCTION

Transit amplification of progenitor cells expands progenitor pools by successive rounds of symmetrical division (Fujita, 1967; Espinosa & Luo, 2008; Legue *et al*., 2015; Legue *et al*., 2016). This allows for rapid assembly of large neural structures in development and is thought to be key for evolutionary adaptation in the development of complex neural circuitry (Borrell & Gotz, 2014). In both the neocortex and cerebellum, elaboration of specialised, transient laminae supporting transit amplification is associated with increased foliation and complexity. Specialised sub-ventricular cell types which are either diminished or absent in mice, are expanded in the human cortex (Hansen *et al*., 2010; Heide & Huttner, 2021) and, as recently shown in human (Haldipur *et al*., 2019), uniquely characterise the progenitor zone of glutamatergic neurons in the cerebellum: the rhombic lip (Wingate & Hatten, 1999).

Like the cortical subventricular zone, the cerebellar EGL is a site of transient, transit amplification, but only for a single cell type: the glutamatergic cerebellar granule neuron. After migrating from the rhombic lip (Wingate & Hatten, 1999; Wingate, 2001; Machold & Fishell, 2005; Wang *et al*., 2005) granule progenitors accumulate within the EGL and undergo multiple rounds of symmetric division (Espinosa & Luo, 2008; Legue *et al*., 2015), driven by Purkinje cell derived Sonic hedgehog (Shh) (Dahmane & Altaba, 1999; Wallace, 1999; Wechsler-Reya & Scott, 1999) before exiting the cell cycle. Granule cells then transition through a range of morphologies in the outer, middle, and inner EGL (Hanzel *et al*., 2019), before undergoing radial migration into the internal granule layer (IGL) and extending characteristic T-shaped axons into a now largely cell body-free molecular layer (Cajal, 1890; 1911; Leto *et al*., 2016; Hanzel *et al*., 2019). Well-characterised molecular cues guide migration of differentiating granule precursors (Hatten & Heintz, 1995; Chedotal, 2010), however regulation of the precise tempo and timing of events is poorly understood.

This current study is prompted by somewhat contradictory observations that urge a clarification of the different roles of Bone Morphogenic Protein (BMP) signalling *in vivo* at the key points of granule cell specification, migration, proliferation, and differentiation. This is important for not only understanding normal cerebellar development but also the origins of medulloblastoma, a devastating childhood brain tumour which traces its cells of origin to both the rhombic lip (Hendrikse *et al*., 2022; Smith *et al*., 2022) and EGL (Millard & De Braganca, 2016).

Firstly, is BMP acutely required for granule cell precursor specification? Early in cerebellar development, a combination of BMP and Delta/Notch signalling is required to specify the rhombic lip (Machold *et al*., 2007; Broom *et al*., 2012), which will then give rise to granule precursors. Correspondingly, exogenous BMP can induce granule cell fate in uncommitted progenitors (Alder *et al*., 1996; Alder *et al*., 1999). However, granule cell specification is not blocked when BMP signalling is attenuated in various transgenic mouse models (Qin *et al*., 2006; Fernandes *et al*., 2012; Tong & Kwan, 2013; Owa *et al*., 2018). Resolving this paradox is important given the recent discovery that the rhombic lip is the origin of the most common Group3 and 4 medulloblastomas (Hendrikse *et al*., 2022; Phoenix, 2022; Smith *et al*., 2022; Williamson *et al*., 2022).

Secondly, what is the role of BMP in EGL assembly and proliferation? Once granule cell progenitors have assembled an EGL, BMP can act to both suppress proliferation and promote differentiation (Zhao *et al*., 2008; Ayrault *et al*., 2010) through its ability to antagonise Shh signalling (Rios *et al*., 2004). Upregulation of Shh-dependent granule cell proliferation results in a larger cerebellum experimentally (Corrales *et al*., 2006) and is associated with a specific group of SHH medulloblastomas (Pietsch *et al*., 1997; Raffel *et al*., 1997; Vorechovsky *et al*., 1997; Dahmane & Altaba, 1999; Wallace, 1999; Wechsler-Reya & Scott, 1999). However, while BMP has therefore been invoked as a potential treatment (Zhao *et al*., 2008; Zhang *et al*., 2011), the same conditional deletions of BMP pathway elements that fail to block early granule cell specification at the rhombic lip do not result in a larger cerebellum as might be expected, but either have no affect (Tong & Kwan, 2013) or generate an EGL that is either smaller (Qin *et al*., 2006; Fernandes *et al*., 2012) or disorganised (Owa *et al*., 2018).

To address these two questions, we designed experiments to precisely manipulate BMP levels during cerebellum development. We show that BMP signalling has distinct and dynamic activity throughout granule cell development in both human and avian models. Experimental inhibition or activation of receptor pathways show that granule cell precursors do not require BMP for their induction, but that their subpial migration to form an EGL and how long they populate the EGL before migrating into the internal granule cell layer is dependent on appropriate BMP signalling. Our human data reveal modifications to the spatiotemporal dynamics of BMP signalling in development that suggest a role in in sustaining the EGL over protracted development of the human cerebellum.

## RESULTS

We first assessed BMP signalling activity during EGL formation and cerebellar foliation in human and chick. We then took advantage of the ability to experimentally manipulate BMP signalling in a targeted manner in the developing avian cerebellum monitoring both the formation of the EGL and an internal granule cell layer.

Changes in BMP signalling in the EGL of the chick correlate with foliation.

Figure 1 shows the comparative timeline of morphological development in the chick, mouse, and human cerebellum (Fig. 1A), and the planes of sectioning used in this study (Fig.1B). For more in depth comparison between mouse and human, we refer the reader to (Haldipur *et al*., 2022). We monitored BMP signalling by assessing the phosphorylation of the highly conserved serine residues (Fig.2A) of Smad proteins using an antibody against the phosphorylated forms of Smads 1, 5, and 9 (collectively; pSmad) (Andrews *et al*., 2017; Owa *et al*., 2018; Najas *et al*., 2020). We confirmed that the residues of Smads 1, 5 and 9 that undergo phosphorylation to activated BMP signalling are highly conserved (Fig.2A). In chick, at embryonic day 5 (E5), pSmad expression is limited to cells proximal to the interface between the neurogenic neuroepithelium and the non-neurogenic roof plate, the rhombic lip (Fig.2B). The expression of pSmad is uniform throughout the EGL during its formation beneath the pial surface of the cerebellum through to E8 (Fig.2C). The expression of pSmad then decreases in the EGL as the cerebellum begins to form folia from E10 (Fig.2D). This is complemented by an increase in expression within the IGL. By E14, expression of pSmad is seen in only a small number of EGL granule cell precursors at the crests of folia (Fig.2E) and is entirely absent from the EGL in the fissures (Fig.2F). A corresponding pattern of maturation is visible in Calbindin-positive Purkinje cells, which display a mature monolayer at the folia troughs and a deep Purkinje cell layer (PCL) that is 3-4 cells deep at the folia crests. The distribution of pSMAD was quantified in the EGL, PCL and IGL (Fig.2G) relative to folia morphology showing that a varying pattern of expression between tips and troughs is shown only in the EGL (Fig.2G) as summarised in Fig.2H. Correspondingly, *in-situ* hybridisation for BMP ligands *Bmp2* and *Bmp4* and receptors *BmpR1a* and *BmpR1b* (Fig.2I) reveals strong expression in the EGL at E10. By E14, BMP expression appears stronger in the folia crests. Similarly, BMP receptors *BmpRIa* and *BmpR1b* also show uniform expression throughout the EGL at E10 but are upregulated within the folia crests at E14.

**Fig. 1.**
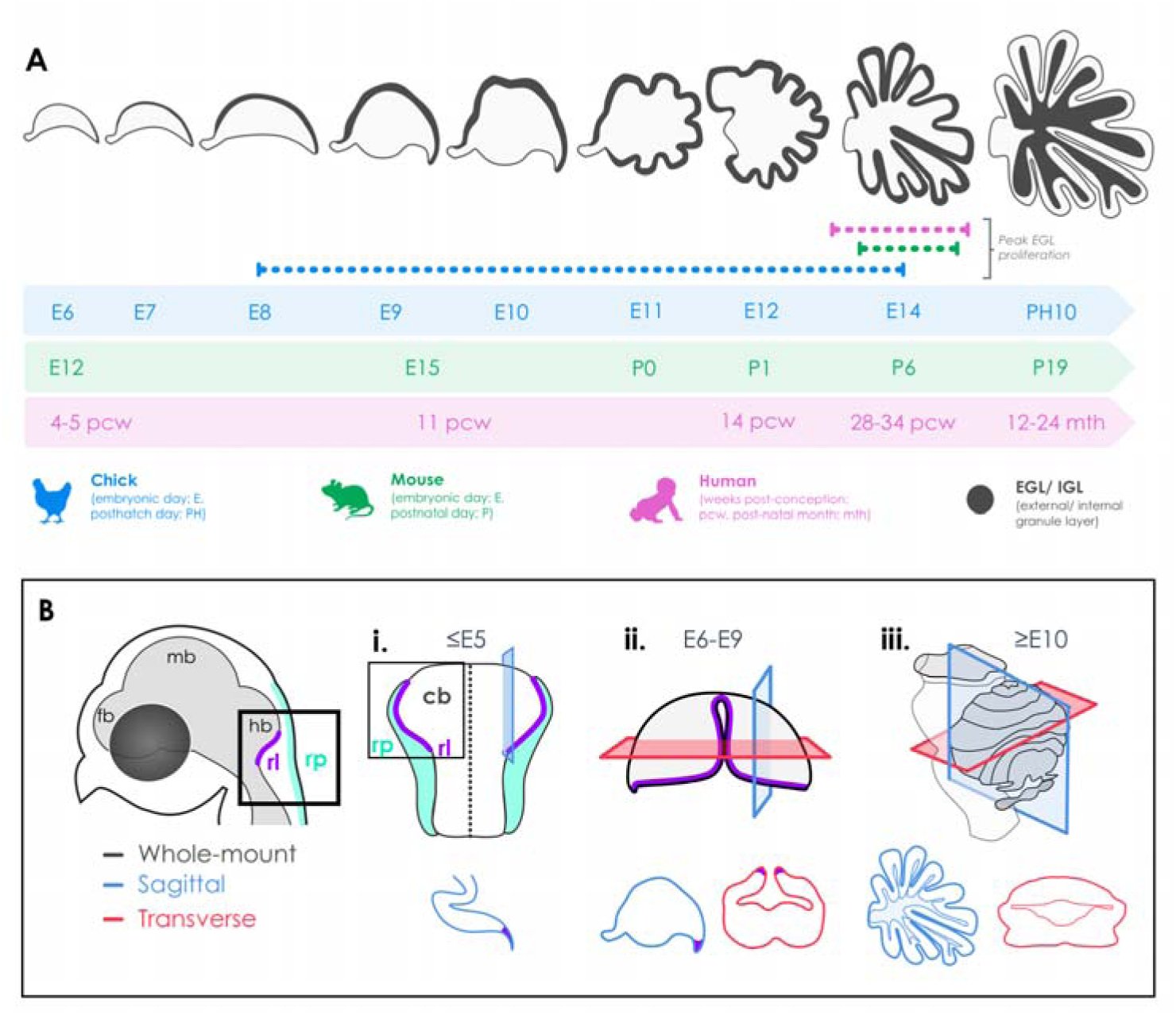
Comparative developmental timescale of the progression of cerebellar foliation in the chick, mouse and human and planes of sections used throughout this study . **(A)** In the chick (blue), granule cell precursors are born from the rhombic lip around embryonic day 6 (E6), forming the EGL by E7, transit amplification ensues from E8 with the onset of foliation around E10 and its completion by 10 days post-hatching (P10). In the mouse (green), granule cells are derived from embryonic day 12 (E12), are proliferative by E15 with foliation beginning around E19 and continuing until postnatal day 19 (P19). In humans (pink), granule cells start to be generated around 9 post-conception weeks (9PCW), with the cerebella foliation starting between 13-17PCW and continuing until around 8 months (8 mth) postnatally. Dotted lines show the timeline of peak EGL proliferation in each species. Schematic not to scale. **(B)** A schematic showing the anatomy of the developing chick hindbrain (rl; rhombic lip, rp; roof plate, fb; forebrain, mb; midbrain and hb; hindbrain), how samples were orientated for whole mount (i) and the planes of section used throughout this study for samples between E6-E9 (ii) and greater than E10 (iii), which were either sagittal (blue) or transverse (red). These symbols are used throughout the figures to orientate the reader.

**Fig. 2:**
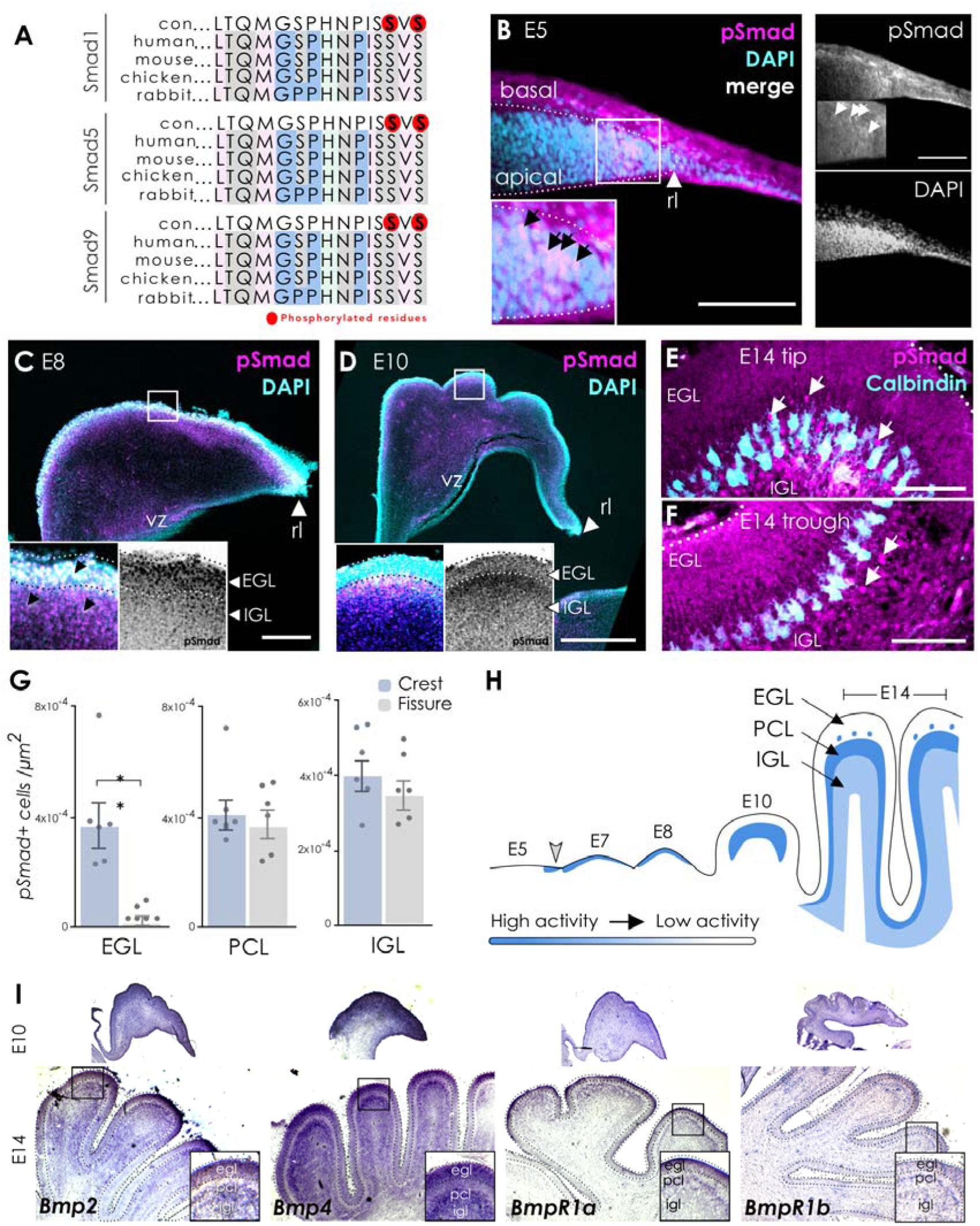
Characterisation of BMP activity and expression throughout the development of the chick cerebellum. An antibody against the conserved phosphorylated 3’ serine residues of Smad1/5/9 (**A**; red-circled residues) show the spatial requirements of BMP activity at the E5 rhombic lip (**B**, n=3), during EGL establishment at E8 (**C**, n=3), at the onset of proliferation in the E10 EGL as folia develop (**D**, n=2) and at highly proliferative and foliated stages of cerebellum development (**E**, **F**; E14, n=6. **(G)** Quantification of pSmad expression in the different layers (EGL; external granule layer, PCL; Purkinje cell layer and IGL; inner granule layer) of the crests (blue bars) and fissures (grey bars) shows that BMP activity is significantly higher in the EGL of the folia crests compared to the fissures (**G**; p= 0.0015; mean ± SEM = 3.521e-005 ± 8.116e-006), however no significant differences in activity were observed between the PCL and IGL (n=6 sections from 4 cerebella). **(H)** A summary of BMP activity during EGL formation and folia development in the chick (E5-E14) **(I)** mRNA expression of BMP ligands (Bmp2, Bmp4) and receptors (BmpR1a, BmpR1b) at E10 (n=3) and E14 (n=4). Scale bars B, E; 25µm, C-D; 50µm.

### BMP signalling in the developing human cerebellum

To see whether this pattern of signalling is conserved, we assessed pSmad expression during corresponding stages of development in the human embryo. At 13pcw of development, when the first folia form (Fig. 3A), pSmad is expressed uniformly throughout the EGL in both the crest (Fig.3B) and trough (Fig.3C) of a folium. Expression is also seen in a deeper layer corresponding to the PCL (Fig.3B’ and C’) and IGL Fig.3B’’ and C’’). At 19 pcw (Fig.3D), the cerebellum has pronounced folia. The expression pSmad remains uniform across the EGL (Fig. 3D-F). The expression of pSmad within the presumptive PCL and IIL is also uniform within cells at crests (Fig.3E) and troughs (Fig.3F) But with a pronounced thickening of all layers at the base of each folium.

**Fig. 3.**
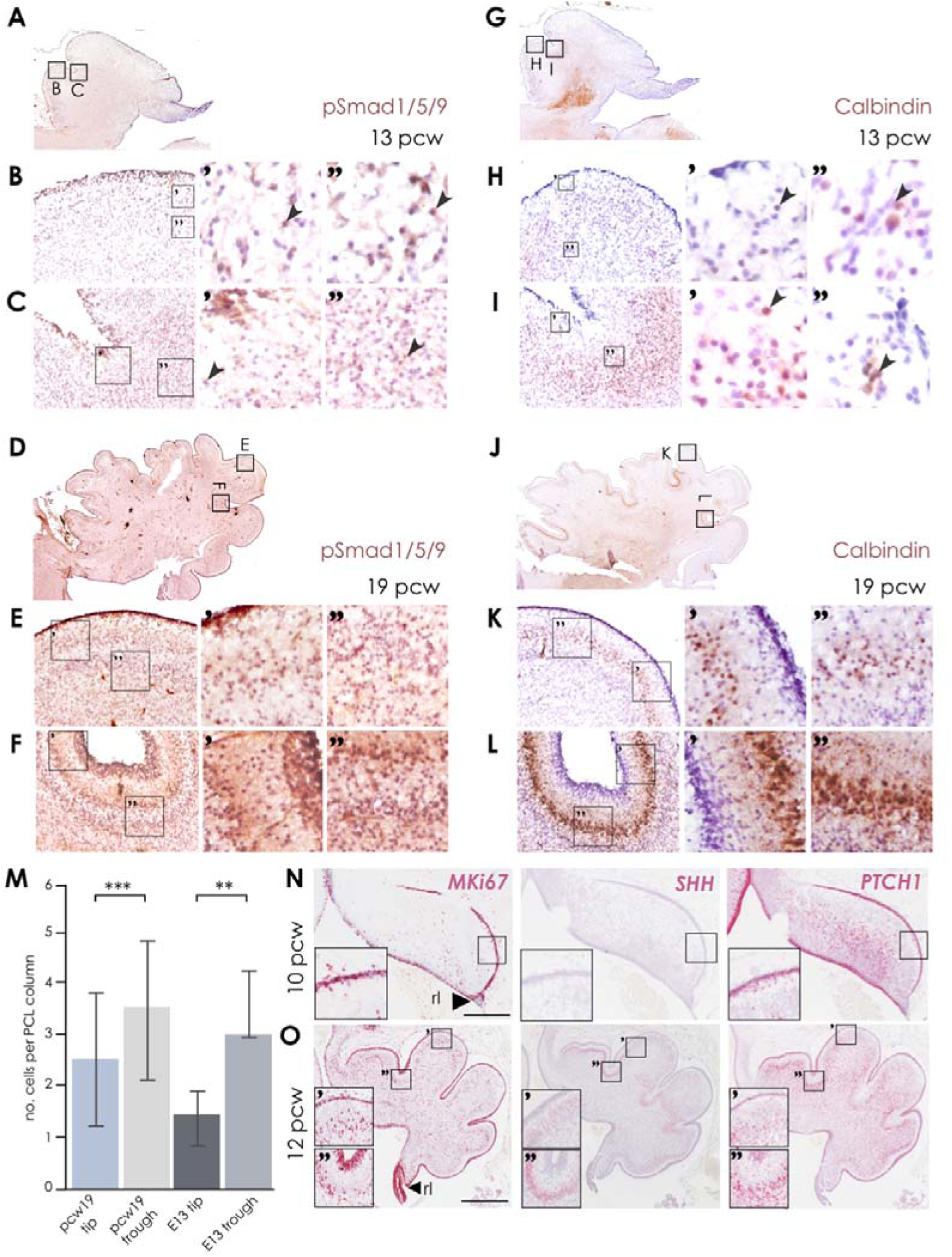
Characterisation of BMP activity, Purkinje cell development and Shh signalling during human cerebellar development. BMP signalling, here determined by pSmad expression, is observed throughout the 13 pcw human cerebellum (A, n=2) folia crests (B) and fissures (C). BMP activity in the 19 pcw human cerebellum (D, n=2) is observed in the EGL of the folia crests (E) and to a lesser extent in the EGL of the fissures (F). Calbindin expression in the 13 pcw cerebellum (D) folia crests (E) and fissures (F) shows the early migration of Purkinje cell precursors. Calbindin expression in the 19 pcw cerebellum (J) shows a less organised Purkinje cell layer in the folia crests (K) when compared to the fissures (L). (M) Quantification of the PCL between tips and troughs of pcw 19 human and E13 chick cerebella. The number of cell soma per dorsal-ventral column were counted. For E13 chick tips, n=85 columns from n=4 sections and troughs n=75 columns from n=4 sections, p=<0.0001; mean ± SEM1.637 ± 0.1486. For human pcw 19, tips n= 79 columns from n=1 section and troughs n=42 from n=1 section, p=0.0002; mean ± SEM 0.9575 ± 0.2483. Expression of the proliferation marker Ki67, Sonic Hedgehog (SHH) and SHH receptor PATCHED1 (PTCH1) during EGL establishment at 10 pcw (N; n=2) and during early foliation at 12 pcw (O; n=1). Scale bars N, O; 500µm.

The differences in layer thickness are mirrored by Calbindin staining for Purkinje cells. At 13pcw (Fig.3G), Purkinje cells are relatively sparsely distributed at a folium crest (Fig.3H) versus trough (Fig.3I). At 19 pcw (Fig.3J), the lower density of Purkinje cells at crest (Fig.3K) versus trough (Fig.3L) is pronounced. The relative numbers of Purkinje cells across folia at each age are quantified in Fig. 3M. As in mouse (Lewis *et al*., 2004; Sudarov & Joyner, 2007) and chick (Dahmane & Altaba, 1999; Rios *et al*., 2004), the initiation of foliation is coincident with the onset of SHH expression in developing Purkinje neurons. An EGL is apparent in the developing human embryo by 10pcw (Fig.3N) as a distinctive sub-pial layer of proliferating cells which express the interphase marker, MKi67. There is no expression of Shh in the PCL although expression of PTCH1 in the EGL indicates that granule cell precursors are transducing a Shh signal, possibly from cerebrospinal fluid. At 12 pcw (Fig. 3O), foliation has commenced and Shh is now strongly expressed in the PCL. Both Purkinje cells and proliferative EGL precursors express PTCH1.

These results show that BMP signalling is occurring in the developing EGL and other cell layers within the cerebellum in both chick and human. BMP signal transduction characterises both Purkinje cells and proliferating granule cell precursors and is sustained in human (Fig.3D-F) compared to chick (Fig.2E). Given its proposed role in antagonising Shh responses, we next chose to investigate how BMPs influence EGL maturation.

### BMP signalling is required for the assembly of the EGL via pial recruitment of granule cell precursors

We investigated the role of BMP signalling in the EGL using electroporation of DNA constructs in chick at the rhombic lip at E4 to selectively target the granule cell precursors. We sought to knock down BMP responses in cells by overexpressing the negative intracellular BMP regulator *Smad6* (Xie *et al*., 2011). By contrast, we aimed to upregulate BMP responses by overexpressing the constitutively active BMP regulator *Smad1EVE*, which is a variant of the transcription factor *Smad1* where the N-terminal SVS residue that is phosphorylated during activation is mutated to EVE (Fuentealba *et al*., 2007; Song *et al*., 2014).

We first confirmed that our constructs were able to affect BMP signal transduction in a predictable manner by characterising the expression of pSmad two days after overexpression at E3 of the control, Smad1EVE, and Smad6 constructs. In all cases, Smad constructs were co-electroporated with a ‘control’ plasmid encoding a fluorescent reporter protein tdTomato or GFP (Fig.4A). Cells were characterised by their expression of GFP and/or pSMAD and the fractions of each quantified following a series of electroporations (Fig.4B). As expected, pSmad expression was either unchanged (control), upregulated in a cell autonomous manner (Smad1EVE) or abolished (Smad6), respectively.

**Fig. 4.**
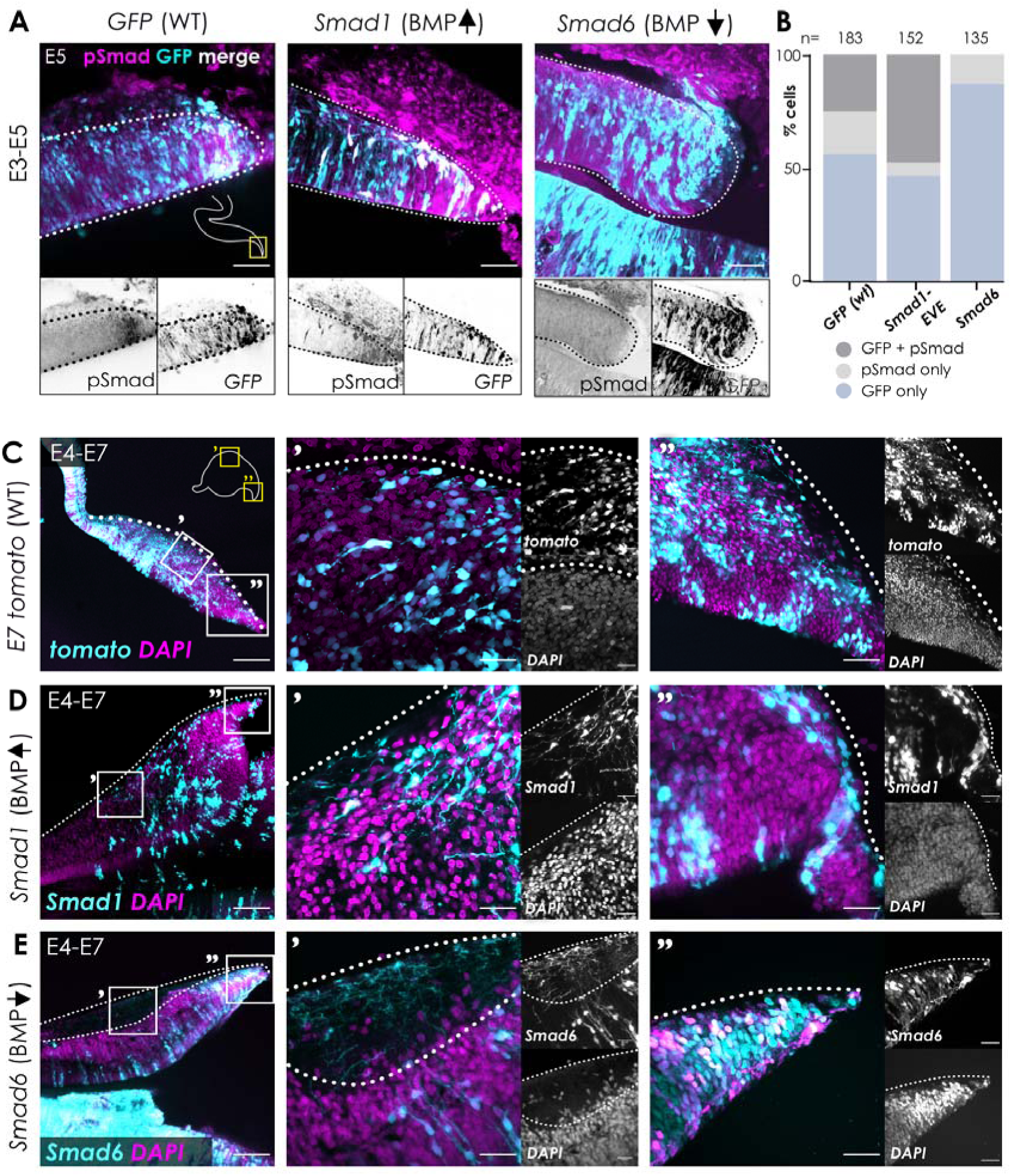
BMP signalling is required for establishing the external granule layer. (A) E3 hindbrains were electroporated with pCAß-GFP alone (n=3), or in combination with pCAß-Smad1EVE-IRES-GFP (Smad1; BMP ; n=3) or pCAß-Smad6 (Smad6; BMP ; n=3), incubated to E5, then embedded in gelatine and sectioned sagittally at 100µm. Sections were immunolabelled using an antibody against phosphorylated Smad1/5/9 (pSmad) to confirm either no change in BMP activity (WT), or up- or down-regulation of BMP signalling at the rhombic lip in Smad1EVE and Smad6 electroporated embryos, respectively. (B) Percentage of cells co-expressing tomato and pSmad following tomato, Smad1EVE or Smad6 electroporation (n= no. cells counted per condition). E4 embryos were electroporated with pCAß-tdTomato alone (C, n=3), or in combination with Smad1 (BMP ; D, n=7) or Smad6 (BMP ; E, n=7), further incubated to E7 and then sagittally sectioned. Dotted white lines A; rhombic lip, C-E; pial surface. Scale bars A; 50µm, C-E; 100µm, C’-E”; 25µm.

Electroporation of a control tdTomato construct into the cerebellar rhombic lip at E4 results in the labelling of the assembling EGL at E7 (Fig.4C). Upregulation of BMP signal transduction by overexpression of Smad1EVE at E4 resulted in tangential migration of EGL cells in a subpial pattern as seen in control electroporations, albeit with a partial depletion of the EGL distal to the rhombic lip (Fig.4D). By contrast, the superficial subpial layer was heavily depleted of all cells (by DAPI label) following inhibition of BMP signal transduction by overexpression of Smad6 (Fig. 4E). This is the location that a granule precursor-rich EGL would be expected to form. The cell free zone was invaded by axonal processes in a manner that is reminiscent of the adult molecular layer and the entire cell-depleted zone was co-extensive with the anteroposterior breadth of the cerebellum (Fig.4E).

### BMP signalling affects the tempo of maturation but not specification of granule cell neurons

To visualise the extent to which disruption of tangential migration from the rhombic lip occurs, we electroporated the rhombic lip at E3 and examined labelled cells in the flat-mounted cerebellum at E5 (Fig. 5A). Manipulation of cell-autonomous BMP signalling did not prevent the formation of rhombic lip derivatives, however their distribution across the cerebellum was altered (Fig.5A) suggesting altered tangential migration. Inhibition of BMP signalling (Fig.5B Smad6; pink line), causes an accumulation of label proximal to the rhombic lip. By contrast, upregulation of BMP (Fig.5B Smad1; yellow line) causes a depletion of tdTomato label close to the rhombic lip (versus control; Fig.5B blue line).

**Fig. 5.**
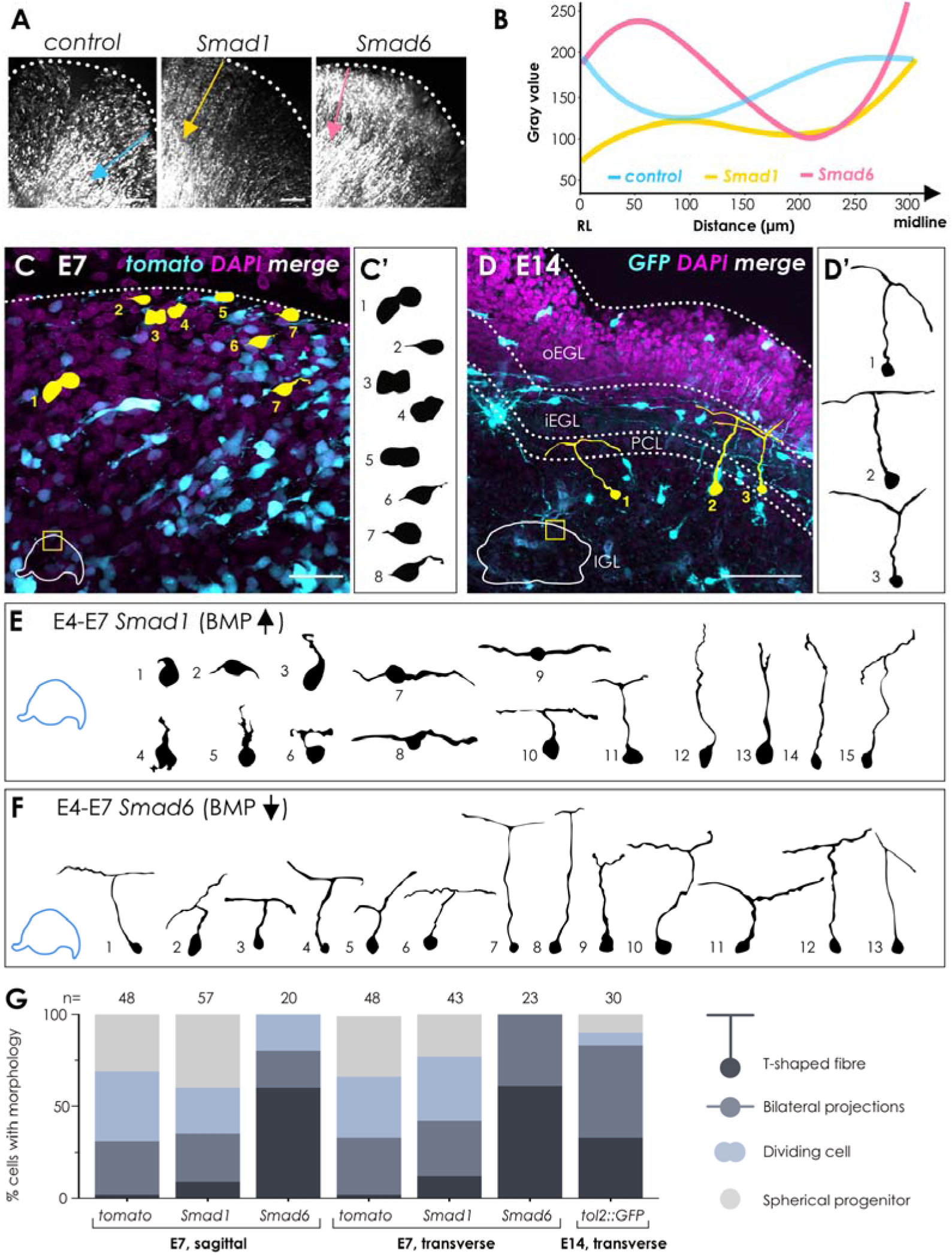
BMP signalling is required for the initial tangential migration of rhombic lip derivatives. (A) Electroporation of tomato, Smad1EVE (Smad1) or Smad6 at the E3 rhombic lip affects migration of early rhombic lip derivatives at E5 (n=3 per condition). (B) The ‘gray’ value plotted against distance from the rhombic lip (coloured lines in A), shows cell density and is indicative of migration from the rhombic lip (n=1 per condition). (C) Sagittal sections from E7 embryos electroporated at E4 with tdTomato control DNA (tomato) show migrating or proliferative granule precursor cells (cell traces C, C’) towards the pial surface (dotted white line). (D) To visualise granule cell morphologies at a foliated stage of development, embryos were electroporated at E2 with tol2::GFP and tol2-transposase and sectioned transversely at E14. Cell traces from the E14 cerebellum show classic T-shaped axon and parallel fibre morphology (D’). (E) Resulting morphologies show varied maturation at E7 following upregulation of BMP signalling at E4, including spherical progenitors (cells 1-6), bilateral progenitors (cells 7-9) and T-shaped axon and fibres (cells 10-15). Mature morphologies are observed at E7 following knock-down of BMP signalling at E4 (F). (G) A summary of morphologies observed in each plane of section for each condition (n= no. cells analysed per condition from n=3 cerebellums per condition, all cells with traceable morphologies were selected from compressed Z-stacks). Scale bars A; 200µm, C, D; 25µm. C’-F cell traces not to scale.

We next assessed the progression of granule cell morphologies as they mature from the EGL and migrate into the IGL by electroporating tdTomato into the rhombic lip at E4 and examining the cerebellum at E7 and E14. At E7, within the EGL, granule cell precursors exhibit a variety of morphologies consistent with both proliferation and tangential migration of bipolar or unipolar precursors (Fig.5C and C’). At this stage, there is no radial migration of granule cells, and the IGL has yet to form. At subsequent stages, the presumed dilution of cellular plasmid concentration through successive rounds of division after electroporation led to a depletion of label within the EGL (data not shown). Therefore, to examine granule cell morphological maturation we used the Tol2 transposon system (Sato *et al*., 2007) to indelibly label granule cell progenitors and their progeny in a mosaic fashion by electroporation of the neural tube at E2. Using this approach, we could visualise mature granule cells with T-shaped axons at E14, within the IGL (Fig.5D and 5D’).

We then asked whether the normal progression of cellular maturation was altered by manipulation of BMP signalling. Overexpression of the constitutively active BMP receptor Smad1EVE within the EGL produced a normal range of cell morphologies at E7 (see Fig.5E’ cells 1-9), interspersed with more mature granule neuron morphologies labelled cells deep to the EGL (Fig.5E cells 10-15), not normally seen at this age (see Fig.5C and C’). This was consistent with a subset of granule cell progenitors differentiating prematurely into IGL neurons. Overexpression of Smad6 at E4, which results in a loss of EGL (Fig. 4E), resulted in a population of uniformly mature granule cells (Fig.5F) at E7 suggesting that cells generated at the rhombic lip (Fig.5A), in the absence of transit amplification, develop prematurely into definitive granule neurons. Furthermore, the normal transverse parallel alignment of axons is perturbed such that normally transversely orientated T-shaped axons are visible in the sagittal sections (Fig. 4E). These results indicate that while altered BMP signalling does not affect the specification of granule cells, it impacts tangential migration, the formation of the EGL, and the timing of differentiation and the precision of axonogenesis, consistent with a role in regulation of the tempo of granule cell maturation.

### Upregulation of BMP signalling accelerates EGL maturation

Since electroporation of Smad1EVE resulted in a partially precocious differentiation of granule cells, we decided to follow the morphological maturation of granule cells in the EGL beyond E7. Between E7 and E8 the EGL shows an increase in proliferation and size (Fig.6A) and Shh signal induction, indicated by *Ptch1* expression in the EGL (Fig.6B). A 4-fold increase EGL thickness (Fig.6C) correlates with significant increase in mitotic marker (PH3) density (Fig.6A). Correspondingly, labelling of the rhombic lip by electroporation at E4 yields cellular morphologies at both E7 (Fig. 5C’) and E8 (Fig.6E, F) that are indicative of proliferative divisions in the outer EGL.

**Figure 6:**
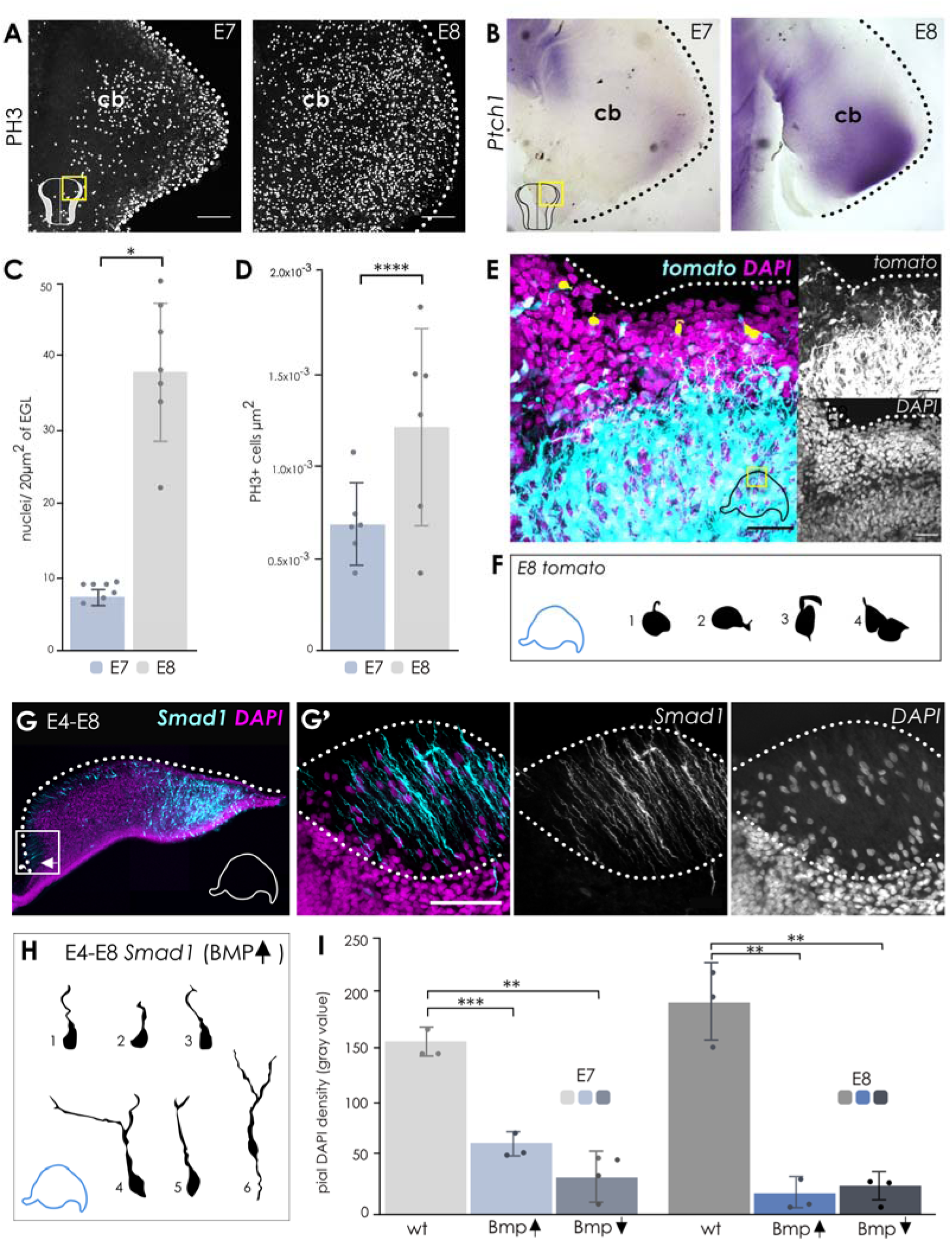
Upregulation of BMP signalling reveals a temporal switch in GCP responsiveness at E8, coinciding with the onset of SHH-induced proliferation. (A) Proliferation, determined by immunolabelling of PH3, at the pial surface greatly increases between E7 and E8, which coincides with the onset of Shh signalling at E8, as determined by expression of Ptch1 mRNA (B). The nuclei density within the EGL (C) significantly increases between E7 and E8, as determined by the number of DAPI-positive nuclei per 20µm^2^ bin of the EGL (n=6 bins per sample from n=7 for E7 and n=7 for E8; p<0.0001, mean ± SEM = 30.37 ± 3.556). Quantification of PH3 immunolabelling (D) of the whole-mounted cerebella (A) shows that there is a significant increase in proliferative activity between E7 and E8 (p=0.0465, mean ± SEM = 0.0005322 ± 0.0002344; n=6 for both E7 and E8). In a control cerebellum electroporated with tdTomato at E4 and sagittally sectioned at E8 (E; n=3), granule cell precursors still exhibit migrating or proliferative morphologies (F). In embryos electroporated at E4 with Smad1 and sectioned sagittally at E8 (G; n=4), an anterior-posterior gradient of EGL loss is observed, with a molecular layer forming in place of the EGL (G’), within which fibres can be seen extending (Smad1, cyan) and sparse DAPI labelling. (H) Granule cells in the E4-Smad1-electroporated cerebellum exhibit mature granule cell morphologies at E8. (I) To show the extent of the depletion of the EGL in control (wt), Smad1 (Bmp ↑) and Smad6 (Bmp ↓) electroporated cerebella, the fluorescent intensity of DAPI labelling at the pial surface was quantified (n=3 samples quantified for each condition at each developmental stage). Significant losses of DAPI density were observed at both E7 and E8 for Smad1 (E7; p=0.0176, mean ± SEM = 92.33 ± 12.41, E8; p=0.0073; mean ± SEM = 173.7 ± 14.97, n=3 per condition) and Smad6 (E7; p=0.0046, mean ± SEM = -124.4 ± 8.446, E8; p=0.0248; mean ± SEM =165.8 ± 26.58, n=3 per condition). Scale bars A; 200µm, E, G’; 25µm. Cell traces in F and H not to scale.

Upregulation of BMP signal transduction at E4 results in accelerated granule cell development and a partial loss of the EGL at E7 (Fig.4D), consistent with the increase in mature cell morphologies observed (Fig.5E). Examining the results of the same manipulation a day later at E8, we find that the EGL is completely depleted of cells (Fig.6G). At high magnification, the EGL has been replaced by a largely cell-free superficial layer containing GFP labelled processes (Fig.6G’). Reconstruction of the morphology of single labelled cells reveals that they bear the hallmarks of mature granule cell morphology: a cell body within the inner granule layer and a T-shaped axon (Fig.5D’). Thus, BMP upregulation results in the depletion of a short-lived EGL to produce a cell-free superficial layer that is reminiscent of an adult molecular layer. The extent of this loss is comparable to that seen when BMP is down regulated in a cell-autonomous manner (Fig.4E), where, by contrast, the formation of an EGL is inhibited. Quantification of EGL cell density at E7 and E8 following electroporation at E4 shows that both upregulation and downregulation of cell autonomous BMP signal transduction results in a similar loss of cell density in EGL at E8 (Fig.6I).

### Granule cells respond to changes in BMP signalling in a cell-autonomous manner

To assess whether the effects of the manipulations of BMP signalling by electroporation are cell autonomous, we devised a strategy to label control and cells with altered BMP signalling side by side in the same embryo. Embryos were electroporated at E2 with tol2::gfp and tol2-transpose, which results in an indelible fluorescent label of cells that take up both plasmids. The same embryos were then electroporated again at E4 at the rhombic lip with either a control plasmid expressing tomato (Fig.7A-C), or a control plasmid in conjunction with the inhibitory Smad6 construct (Fig.7D). This approach was expected to label a large cohort of rhombic lip derived cells that were electroporated at E2 with the control tol2-gfp construct, but only a subset of later-targeted (E4) rhombic lip derived cells due to the expansion in size of the cerebellum. Accordingly, at E7 in sagittal section, whereas E2 GFP-labelled cells fill the cerebellum, E4 tomato co-labelled migratory rhombic lip derivatives are restricted to sub-pial stream (Fig.7B), within the nuclear transitory zone (Fig.7B’) and within the nascent EGL (Fig.7B’’). In the latter, tangentially migrating EGL cells are visible that are either labelled only by the e2 electroporation (Fig.7C, left arrow), or else double labelled (Fig.7C, right arrow). To assess the effect of BMP signal downregulation, a second group of embryos were electroporated at E4 at the rhombic lip with a combination of tomato and Smad6. This allowed us to distinguish, in sagittal section, cell autonomous and non-autonomous effects of the inhibition of BMP signal transduction (Fig.7D). Cells expressing GFP only (Fig.7D’) are distributed throughout cerebellar layers including the superficial, sub-pial migratory layer. By contrast, cells expressing tomato (and hence Smad6) were excluded from this layer (Fig.7D’’) Reconstruction of cells labelled with GFP only in the sub-pial layer showed unipolar or bipolar cell morphologies consistent with tangential migration (Fig.7D’: red, green, and purple arrows). Cells expressing both GFP and tomato (and hence Smad6) displayed T-shaped axons that are characteristic of mature granule cell morphology consistent with those induced where Smad6 only was electroporated (Fig. 5F).

**Fig. 7.**
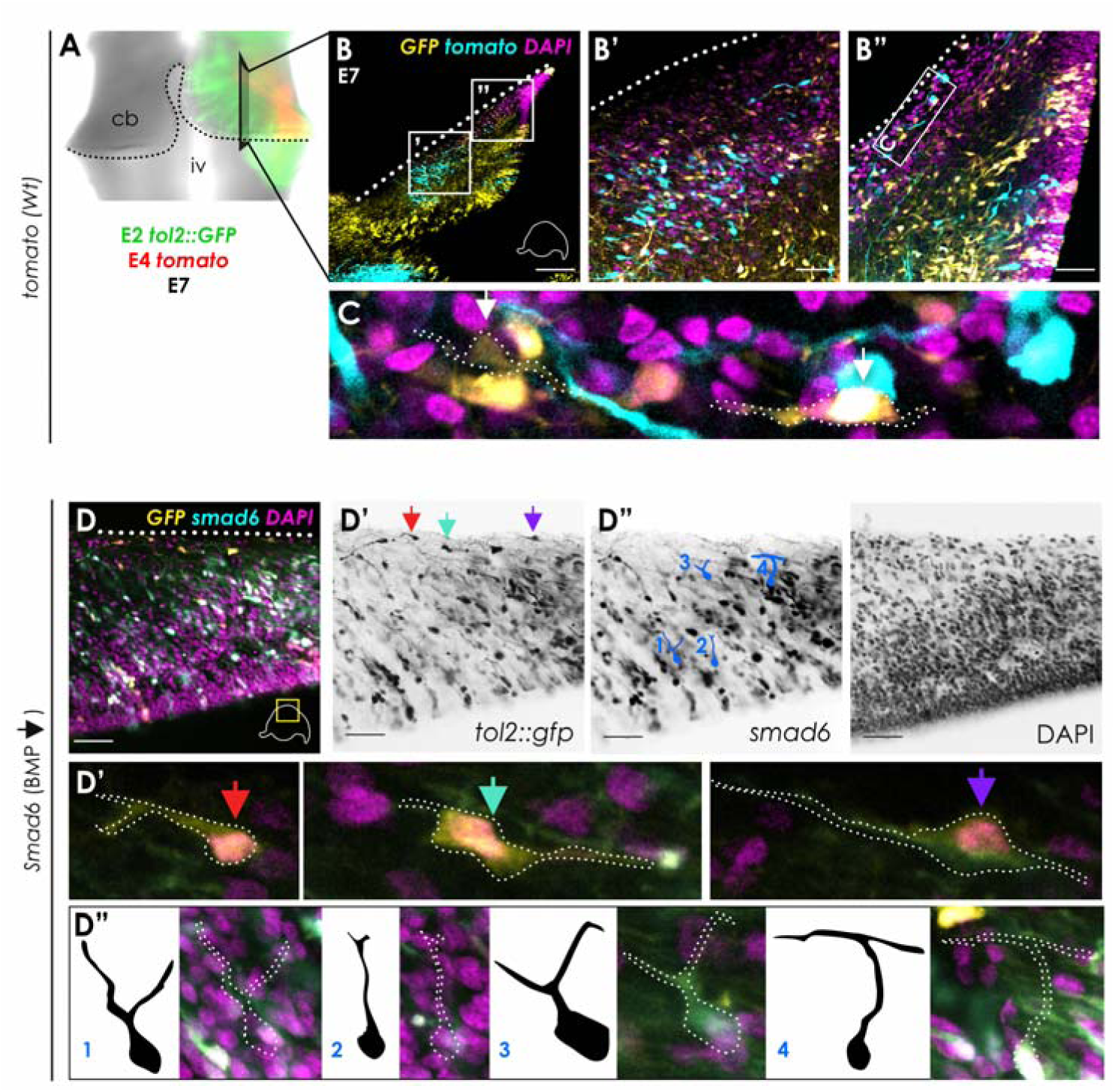
Following BMP-manipulated cell behaviour in a tol2::gfp wildtype background. E2 embryos were electroporated with tol2::gfp and tol2-transposase in order to permanently label the progenitors and their progeny within the developing cerebellum with tol-2::gfp. The same embryos were then electroporated for a second time at E4 with either tomato alone (A-C) or in combination with Smad6 (D) or Smad1EVE (not shown) and incubated until E7, whereby both electroporations can be visualised on the intact cerebellum (A). In sections from cerebella electroporated with tomato only at E4, rhombic lip derivatives expression both tol2-gfp and tomato are observed migrating across the pial surface (B”). In cerebella electroporated with Smad6 at E4, cells expressing gfp only are observed migrating along the pial surface (D’, coloured arrows), whereas cells expressing both tomato and smad6 migrate ventrally following their specification from the rhombic lip, and prematurely differentiate (D’, cells 1-4).

## DISCUSSION

In this study we examined the phosphorylation of SMAD during cerebellar development of chick and human revealing contrasting patterns of BMP signalling in the EGL. These findings are summarised in Figure 8. In chick, the cell-autonomous manipulation of BMP signalling revealed that BMP is required for the sub-pial migration of granule cells and regulates the tempo of maturation in the EGL. Downregulation of BMP signal transduction drives granule cells directly into the internal granule cell layer. Upregulation of BMP accelerates the maturation of granule cells in the EGL. Both manipulations thus result in the precocious appearance of post-mitotic granule cell T-shaped axons.\

**Fig. 8:**
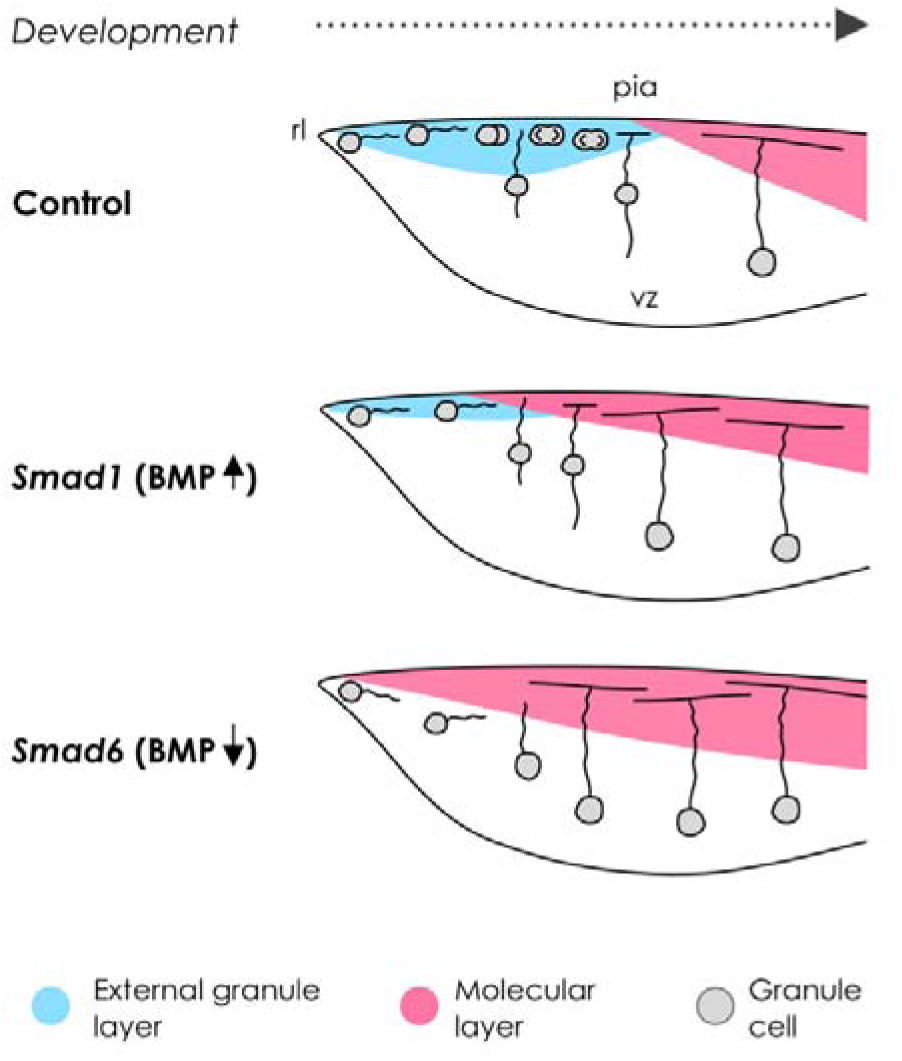
Tightly regulated spatiotemporal activity of the BMP signalling pathway is required for proper development of the external granule layer. In control cerebella, granule cell precursors (grey circles) migrate tangentially from the rhombic lip (rl) across the pial surface to form the external granule layer (blue), where they undergo massive transit amplification before they exit the cell cycle, differentiate, and migrate radially towards the inner granule layer, eventually leaving behind a molecular layer (pink) in which their parallel fibres extend. In cerebella with increased levels of BMP signalling (Smad1, BMP ↑), granule cells are initially recruited to the pial surface, however coinciding with the onset of Shh at E8 (in chick), the excess of BMP signalling forces granule cells to exit the cell cycle prematurely and migrate, leaving behind a molecular layer and a depleted cerebellum. Following inhibition of BMP signalling at the rhombic lip (Smad6, BMP ↓) cells fail to be recruited to the pial surface, and instead immediately migrate ventrally and radially, before extending parallel fibres in a disorganised arrangement into the molecular layer that forms at the pial surface.

### Granule cell production is independent of both BMP signalling at the rhombic lip

Our observation that granule cell precursors are produced at the rhombic lip regardless of cell autonomous disruption of BMP signalling appears to contradict the prevailing model of cerebellar development: that BMP secreted by the roof plate cells induces rhombic lip. BMP is not only appropriately expressed in cells adjacent to the rhombic lip (Campo-Paysaa et al., 2019), but addition of BMP to a culture of cells from embryonic rostral hindbrain can also induce granule cell formation (Alder *et al*., 1999). This apparent discrepancy can be resolved in a model where BMP signalling is required at early stages to dorsalise the neural tube (Alder *et al*., 1999), but not acutely to produce rhombic lip derivatives. By contrast, induction of rhombic lip derivatives relies on acute notch mediated cell-cell interactions (Machold *et al*., 2007; Broom *et al*., 2012) between neural progenitors and closely apposed non-neural cells which express BMP, at the edge of the roof plate (Campo-Paysaa *et al*., 2019). Thus, rhombic lip derivatives can be generated in conditional mouse mutants that interrupt the BMP pathway (Qin *et al*., 2006; Fernandes *et al*., 2012; Tong & Kwan, 2013) but not in conditions where the early dorsalisation of the neural tube is blocked.

### Granule cell production can be independent of the formation of an EGL

It has long been presumed that granule cell morphology Is a product of the intricate step-wise developmental choreography within the EGL described by Cajal (Cajal, 1894). In this scheme, T-shape axons are formed as cells first exit the cell cycle in the EGL and then both extend and anchor postmitotic granule neurons as they migrate radially into the IGL. However, the presumed, obligate sequential steps of granule cell maturation have been questioned by recent studies. Not only are the early sequence of events in maturation occasionally reversed in chick (Hanzel *et al*., 2019), but the granule cell layer can be replenished in the absence of an EGL in regenerating mouse cerebellum (Wojcinski *et al*., 2017). Similarly, our results show that in conditions of attenuated BMP signalling, post-mitotic granule cells develop directly from the rhombic lip, bypassing the formation of a transient amplifying layer (Fig.8).This is identical to the effects of overexpression of NeuroD1 at the rhombic lip (Butts *et al*., 2014; Hanzel *et al*., 2019) and mimics the direct development of granule cells in fish, which lack an EGL (Gona, 1976; Chaplin *et al*., 2010; Butts *et al*., 2014). These collected observations argue that the EGL is not necessary for granule cell production but is rather an adaptation related to efficiently organising proliferation (both expanding the number and duration of granule cell production) and dispersal of granule cell during the development of amniote cerebella (Chaplin *et al*., 2010).

### BMP signalling regulates the tempo of granule cell maturation in the EGL

Whereas the down regulation of BMP signal transduction suppresses EGL formation, constitutive upregulation appears to shorten the duration of granule cell transit amplification (Fig.8), such that the EGL is rapidly depleted of cells by E8. The resulting cell free layer is filled with granule cell axons, consistent with the precocious transition of the EGL into a molecular layer that normally appears at E15 (Bastianelli, 2003). This corresponds with *in vitro* evidence showing that, in culture, terminal divisions are promoted in granule cell precursors when proliferation rates are upregulated by Shh in a background of constitutively raised BMP signal transduction (Rios *et al*., 2004; Zhao *et al*., 2008). These observations are consistent with a model whereby BMP promotes neurogenic divisions at the expense of self-renewing, transit amplification divisions that characterise the EGL (Nakashima *et al*., 2015; Yang *et al*., 2015), reviewed by (Le Dreau, 2021). A similar role for BMP signalling appears to drive terminal differentiation and radial migration of upper layer cortical progenitors (Saxena *et al*., 2018) . As in the cerebellum, the proliferation of transit amplifying cortical glutamatergic progenitors in the cortex is also enhanced in conditions of raised Shh (Wang *et al*., 2016). Thus, BMP antagonism of Shh-driven proliferation may be a general mechanism for regulating terminal differentiation in large neuronal populations in the amniote brain.

### BMP signalling as a regulator of the lifespan of the transient EGL

Human cerebellum development is notable for the extremely extended duration of transient amplification in the EGL that lasts over a period of months from 30 days post-conception until two years after birth (van Essen *et al*., 2020). This is supported developmentally by the adaptation of the rhombic lip to adopt a subventricular zone (SVZ) that presumably facilitates an extended production of rhombic lip derived progenitors (Haldipur *et al*., 2019), and alteration of transcription factor expression that extends their developmental lifespan (Behesti *et al*., 2021). Our results show that BMP signalling in the EGL of the human embryo shows a remarkably uniform distribution that is independent of morphogenic patterns of foliation that correlate to BMP signal variation in the developing chick cerebellum. In the chick, uniform BMP signalling is a hallmark of the early developing EGL. By contrast, in the mouse, BMP upregulation corresponds to the disappearance of the transient EGL (Owa *et al*., 2018). These collected observations suggest that it is the balance of BMP signalling against a background of proliferation that modulates the rates of areal expansion (in foliation) or extinction of the EGL, presumably through regulating the balance of self-renewing versus terminal divisions. In humans, a uniform BMP expression speaks to regulation of a signal that maintains proliferation throughout morphogenic foliation without triggering a depletion of the EGL.

## Conclusions

Overall, our results show a central role for BMPs in the formation of the EGL but also show how a balance of BMP signalling regulates the balance of the neurogenic versus self-renewing divisions within the EGL. BMPs are not required acutely for granule cell specification per se, but the scale of granule cell production is intimately dependent on the role of BMP in both forming and then maintaining the transient, transit amplifying EGL. This has implications for understanding the origin and possible treatments of medulloblastoma, where exogenous BMPs that might be effective against SHH-type would not ameliorate the predominant Group 3 and 4 rhombic-lip-derived tumours. In normal development, sustained BMP signalling within the EGL may play an important role in sustaining transit amplification over the protracted developmental time course of the human cerebellum.

## MATERIALS AND METHODS

### In-ovo **electroporation**

Fertilised hen’s eggs (Henry Stewart) were incubated at 38°C at 70% humidity. Electroporations were performed between stages HH10-25 (Hamburger & Hamilton, 1993), or between embryonic day 2 (E2) to E4. Eggs were windowed using sharp surgical scissors and the vitelline membrane covering the head removed. DNA was injected into the fourth ventricle at a final concentration of 1-3 µg/µl in addition to trace amounts of fast-green dye (Sigma). Three 50ms square waveform electrical pulses at 5V (E2) or 10V (≥E3) were passed between electrodes that were placed on either side of the hindbrain Figure 3a). Five drops of Tyrode’s solution containing penicillin and streptomycin (Sigma) was administered on top of the yolk before being resealed and further incubated for the designated number of days. Embryos were fixed in 4% PFA in PBS for 1 hour at room temperature or overnight at 4°C and then processed for histology. Table 1 summarises the DNA plasmid constructs used throughout this study.

**Table 1:** DNA plasmids.

| Plasmid | Details | Source/ reference(s) |
| --- | --- | --- |
| *pCAGGS-Smad6* | BMP inhibitor | Andrea Streit, KCL (Xie et al., 2011) |
| *pCAGGS-Smad1EVE-IRES-GFP* | Constitutively activated Smad1 | Andrea Münsterberg, UEA (Fuentelba et al., 2007; Song et al., 2014) |
| *pCAGGS-GFP* | GFP reporter | Richard Wingate, KCL |
| *pCAGGS-tdTomato* | Tomato reporter |  |
| *pTol2-GFP* | Tol2 integrated GFP reporter | Nico Daudet, UCL (Sato et al., 2007; Freeman et al., 2012) |
| *pTol2-Transposase* | Tol2 transposase enzyme |  |

### Human foetal tissue procurement

Histological analysis of the human cerebellum: Human cerebellar samples used in this study were collected in strict accordance with legal ethical guidelines and approved institutional review board protocols at Seattle Children’s Research Institute, University College London. and N,wcastle University. Samples were collected at by the Human Developmental Biology Resource (HDBR), United Kingdom, with previous patient consent. Samples were staged using foot length with the age listed as post-conception weeks (pcw), which starts from the point at which fertilization occurred.

Samples were fixed in 4% PFA and then processed through alcohol gradients and xylene. Processed tissue was then embedded in paraffin wax prior to sectioning. Samples sectioned using the cryostat were treated with 30% sucrose following fixation. Paraffin and cryo-sections were collected at 4 and 12 μm respectively. In situ hybridization assays were run using commercially available probes from Advanced Cell Diagnostics, Inc. Manufacturer recommended protocols were used without modification. The following probes were used in the study SHH (#600951), MKI67 (#591771) and PTCH1 (#405781). Sections were counterstained with fast green. Images were captured at 20X magnification using a Nanozoomer Digital Pathology slide scanner (Hamamatsu; Bridgewater, New Jersey).

### Tissue processing, immunohistochemistry, **in situ** hybridisation and imaging

Cerebella were dissected between E5-E14 and either whole-mounted in glycerol or embedded in 20% gelatine, 4% low-melting point (LMP) agarose or OCT and sectioned at 50µm using a vibratome (Leica) or at 15µm using a cryostat (Microm). For immunolabelling, whole-mount and gelatine sections were washed with PxDA (1x PBS, 0.1% Tween-20, 5% DMSO, 0.02% NaN3), then 3x 30 minutes, blocked (PxDA, 10% goat serum) 2x 1 hour, and incubated in primary antibody (diluted in block) for 48 hours at 4°C on a rocker. Tissue was washed in block for 5 mins then 3x 1 hour. Secondary Alexaflour (Thermofisher) antibodies were diluted in block (1:500) and incubated overnight at 4°C. Samples were washed 3x 1 hour with block, 3x 3 mins with PxDA and 1 hour in 4% PFA. Sections were mounted using Fluoroshield containing DAPI (Abcam). Frozen sections were thawed at room temperature for 1 hour, washed in 1x TBS buffer (2% BSA, 1x TBS, 0.02% NaN3, pH7.6), blocked in 1x TBS buffer for 10 minutes, and incubated in primary (diluted in 1x TBS) overnight at room temperate in a humidity chamber. Slides were washed in 500ml 1x TBS for 10 minutes (with stirring), then incubated in secondary antibody (biotinylated for DAB staining (1:300) or Alexaflour for fluorescence (1:500; diluted in 1x TBS) for 1 hour at room temperature. For DAB staining, The Strept ABC-HRP (1:100 of each A and B in 1x DAB developing buffer) was left to conjugate for 30 mins. Slides were rinsed in 1x TBS and then incubated in the conjugated Strept ABC-HRP solution for 30 minutes. Slides were rinsed in 500ml 1x TBS for 5 minutes, with stirring, and then developed for 10 minutes in DAB solution (DAKO DAB enhancer was used for pSmad1/5/9 at 1:300). Slides were washed under running water, counterstained with haematoxylin, and returned to the running water until nuclei turned blue. Antibodies, and the dilutions they were used at is summarised in Table 2.

**Table 2:** antibodies used in this study.

| Target | Host species | Dilution | Source |
| --- | --- | --- | --- |
| Phosphorylated Smad1/5/9 (pSmad) | Rabbit | 1:100 | CST 13820 |
| Calbindin | Mouse | 1:1000-5000 | Swant D28K |
| Phosphohistone-3 (PH3) | Rabbit | 1:100 | CST |
| DAPI | n/a | n/a | Abcam |
| Alexafluor-488 | Goat anti-mouse | 1:500 | Thermofisher |
| Alexafluor-488 | Goat anti-rabbit | 1:500 | Thermofisher |
| Alexafluor-568 | Goat anti-mouse | 1:500 | Thermofisher |
| Alexafluor-568 | Goat anti-rabbit | 1:500 | Thermofisher |

For *in situ* hybridisation, dissected hindbrains were fixed in 4% PFA for 1 hour (and stored up to 3 months) and stained as previously described (Myat et al., 1996) using a digoxygenin-labelled riboprobe (Roche) against the target mRNA sequence (Table 3). Tissue was flat mounted in 100% glycerol and imaged from the dorsal side.

**Table 3:** mRNA riboprobes used in this study.

| mRNA target | Sequence accession number |
| --- | --- |
| *Gallus bone morphogenetic protein 2 (BMP2)* | NM_204358.1 |
| *Gallus bone morphogenetic protein 4 (BMP4)* | NM_205237.4 |
| *Gallus bone morphogenetic protein receptor type 1A (BMPR1A)* | NM_205357.2 |
| *Gallus bone morphogenetic protein receptor type 1B (BMPR1B)* | NM_205132.1 |
| *Gallus patched 1 (PTCH1)* | NM_204960.3 |

### Image analysis

Sections with fluorescent labelling were imaged using a Zeiss LSM 800 confocal microscope and Z-projections compiled with ImageJ (Schneider et al., 2012). Non-fluorescent samples were imaged using a Zeiss Axioscope microscope. To represent the pial migration from the rhombic lip (Figure 5a, b) the fluorescence intensity, termed “gray value” in ImageJ, from the rhombic lip towards the midline in an area of abundant electroporation (coloured lines; Figure 5a), was plotted as a surface histogram, obtained from the plot profile plugin and a curve of best fit (5th degree polynominal). ImageJ was also used to quantify the number of antigen-expressing cells per area (+ve cells/μm^2^ ); cells positive for pSmad labelling were manually counted using the cell counter plugin (Figure 4a) whereas quantification of PH3 labelling (Figure 6a,d) was done automatically by converting a compressed Z-stack to a binary image, watershed function applied and the analyse particles plugin applied to count positive cells in the sample. The area of the tissue being quantified was also measured in ImageJ and the number of +ve positive nuclei per µm^2^ was then calculated in Excel and analysed for significance in GraphPad Prism. To measure the density of DAPI +ve nuclei at the pial surface in electroporated samples (Figure 6i), individual slices from Z-stacks of each sample were processed to binary images, and a line was drawn across the pia in ImageJ and the fluorescent density averaged across this line using the plot profile plugin. To analyse the maturity of the PCL, the number of cells in each dorsal-ventral column of the PCL in the folia vs. troughs were averaged across n=4 (E13 chick) and n=1 (pcw19 human) sections.

### Statistical analyses

All data were analysed in GraphPad Prism, and non-paired parametric t-tests were carried out to identify significance.

## Data and Materials availability

The human material was provided by the Joint MRC/Wellcome (MR/R006237/1) Human Developmental Biology Resource (www.hdbr.org). Human tissue used in this study was covered by a material transfer agreement between SCRI and HDBR. Samples may be requested directly from the HDBR.

## Acknowledgments

This work was funded by Queen Mary University, London. We thank Andrea Munsterberg and Grant Wheeler (University of East Anglia) for the *Smad1EVE* construct, Koichi Kawakami (National Institute of Genetics, Japan) and Yoshiko Takahashi (Kyoto University) for the Tol2 construct, and Andrea Streit (King’s College, London) for the *Smad6* construct.

